# PUFFFIN: A novel, ultra-bright, customisable, single-plasmid system for labelling cell neighbourhoods

**DOI:** 10.1101/2023.09.06.556381

**Authors:** Tamina Lebek, Mattias Malaguti, Alistair Elfick, Sally Lowell

## Abstract

Cell communication orchestrates development, homeostasis, and disease. Understanding these processes depends on effective methods to study how cells influence their neighbours. Here, we present Positive Ultra-bright Fluorescent Fusion For Identifying Neighbours (PUFFFIN), a novel cell neighbour-labelling system based upon secretion and uptake of a positively supercharged fluorescent protein s36GFP. To achieve sensitive labelling, we amplified the fluorescent signal by fusing s36GFP to the ultra-bright protein mNeonGreen. Secretor cells transfer s36GFP-mNeonGreen to neighbours while retaining a nuclear mCherry, making it straightforward to unambiguously identify, isolate, and profile neighbours of secretor cells. Transfer of the s36GFP-mNeonGreen fusion occurs within minutes, enabling even relatively transient interactions to be captured or tracked over time with live-imaging or flow cytometry. To enhance flexibility of use, we incorporated HaloTag technology to facilitate colour- of-choice labelling. To further increase the range of applications, we engineered a customisable single-plasmid construct composed of interchangeable components with option to incorporate any additional transgene. This allows users to manipulate the properties of a cell, while at the same time applying a fluorescent label to the surrounding cells. PUFFFIN offers a simple, sensitive, customisable approach to profile non-cell-autonomous responses to natural or induced changes in cell identity or behaviour in mammalian cells.

**Graphical abstract:** 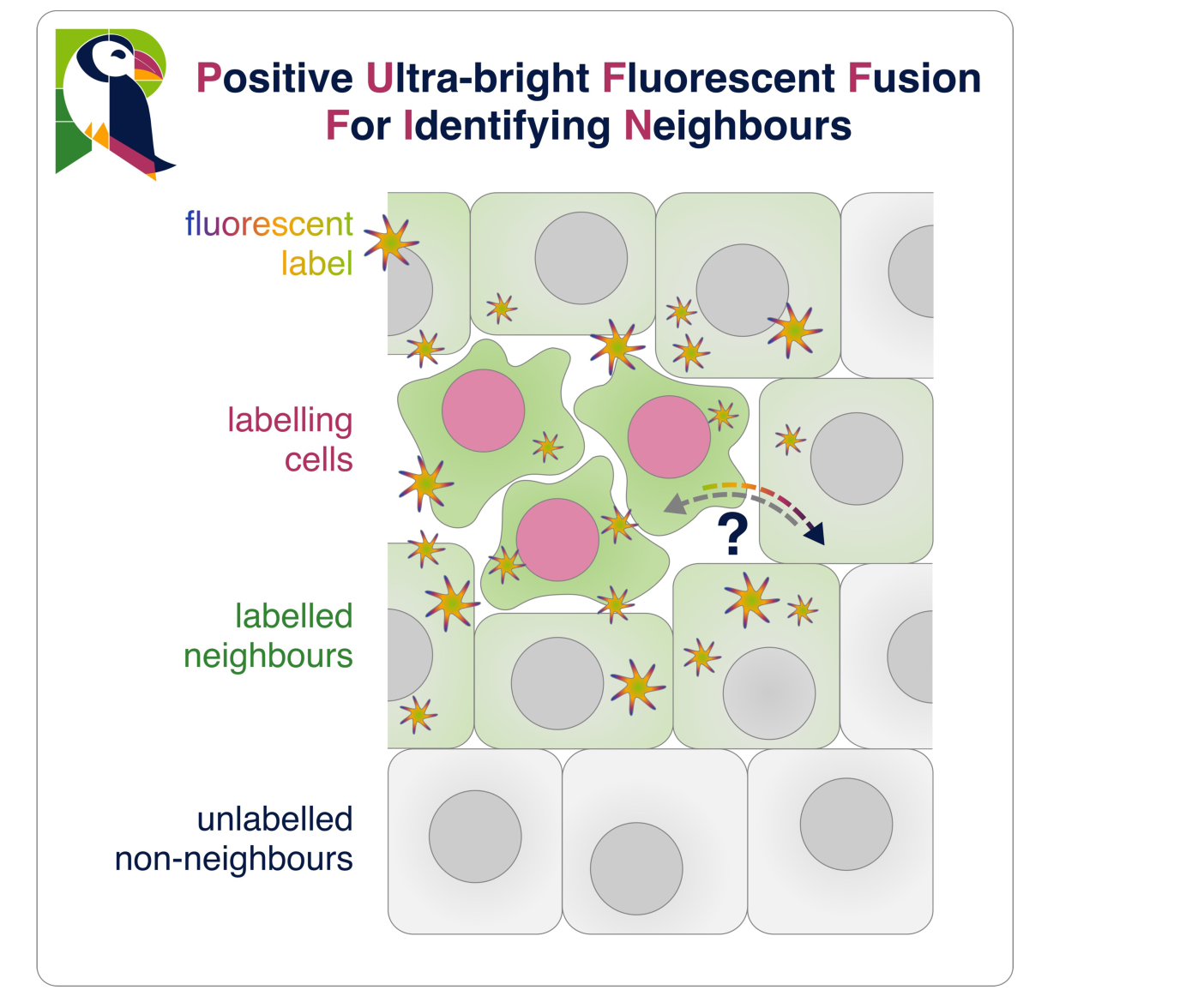

## Introduction

All multicellular life depends on cell-cell communication. It is therefore important to understand the mechanisms by which cells influence their neighbours. Traditional approaches to characterise neighbour responses are often based on image analysis, which is limited by the available combination of fluorescence channels (Lin, Fallahi-Sichani, and Sorger 2015; Lun and Bodenmiller 2020). This limitation is being overcome by advances in spatial transcriptomics technologies (Walker et al. 2022), but these are technically challenging for analysing complex tissues at single cell resolution and remain beyond the technical and financial reach of many laboratories. Alternative approaches are needed to facilitate simple, reliable, sensitive, unbiased profiling of neighbour responses.

One such approach is to engineer synthetic signalling pathways that allow particular cells to induce expression of fluorescent proteins in neighbours, allowing those neighbours to be isolated by fluorescence activated cell sorting (FACS) and then profiled, for example using omics analysis. Recent advances in protein design and cell engineering provide a range of such neighbour-labelling systems with a notable example being synNotch (Gordon et al. 2015; Morsut et al. 2016; Roybal et al. 2016; T.-H. Huang, Velho, and Lois 2016) and its derivatives, such as synNQ (He, Huang, and Perrimon 2017), TRACT (T. Huang et al. 2017), SyNPL (Malaguti et al. 2022), gLCCC (S. Zhang et al. 2022) or diffusibile synNotch (Toda et al. 2020). Other synthetic signalling systems include Tango (Talay et al. 2017; Sorkaç et al., n.d.) and its light-inducible versions (M. W. Kim et al. 2017; C. K. Kim et al. 2019; Cho et al. 2022), G-baToN (R. Tang et al. 2020), and the proximity labelling systems FAP-DAPA (Carpenter et al. 2020), LIPSTIC (Pasqual et al. 2018) and uLIPSTIC (Nakandakari-Higa et al. 2023). One limitation of these powerful systems is that they are based upon synthetic receptor-ligand interactions, so they require genetic modification of ‘sender’ cells and ‘responding’ cells (Morsut et al. 2016; Toda et al. 2020; R. Tang et al. 2020; Malaguti et al. 2022; S. Zhang et al. 2022).

Simpler systems engineer fluorescent molecules for both export by senders and uptake by other cells in the niche. This is made possible by the fusion of signal peptides and cell penetrating peptides, such as the HIV-1 transactivator of transcription (TATk), with fluorescent proteins. This concept has been exemplified by the PTD-GFP system for green fluorescent protein (GFP) (Flinterman et al. 2009) and by the Cherry-niche for the red-fluorescent mCherry (Ombrato et al. 2019; 2021). The Cherry-niche system, in particular, has proven its efficacy in characterising the tumour niche (Ombrato et al. 2019; Nolan et al. 2022) and has the potential for broad applicability for neighbour labelling. Nonetheless, access to alternative systems for secretion and uptake of fluorescent proteins would enhance flexibility in the design of neighbour-labelling experiments.

Supercharged GFP has previously been used as a macromolecule-delivery system (McNaughton et al. 2009; Cronican et al. 2010; Thompson, Cronican, and Liu 2012). It is engineered to have a theoretical net charge of +36 (+36GFP) (Lawrence, Phillips, and Liu 2007), facilitating interaction with cell membranes and subsequent endocytosis (McNaughton et al. 2009; Thompson et al. 2012). +36GFP is efficiently taken-up by mammalian cells (McNaughton et al. 2009), and so proteins fused to +36GFP can be delivered into cells, outperforming fusions to commonly used cell-penetrating peptides (such as the TATk peptide and the Penetratin peptide) in several cell lines (Cronican et al. 2010).

In this study we set out to examine whether +36GFP could be re-purposed for neighbour labelling. To maximise the range of applications, we explored strategies for signal amplification, with the aim of developing a sensitive system that could unambiguously identify even relatively transient interactions and be amenable to live imaging. We also sought simple options for changing the colour of labelling so that the system could be combined with any existing fluorescent reporters, and a customisable design of exchangeable modules that would maximise flexibility of experimental design.

## Results and discussion

### PUFFFIN can unambiguously label surrounding cells

We set out to design a system that could facilitate unambiguous labelling of neighbours after delivery of a single plasmid (**Figure 1A**). The supercharged green-fluorescent protein +36GFP (**Figure 1B**), is a positively-charged version of GFP (Lawrence, Phillips, and Liu 2007) that can be taken up by mammalian cells when exogenously introduced to cultures (McNaughton et al. 2009; Cronican et al. 2010; Thompson et al. 2012). We therefore decided to assess whether +36GFP could form a core component of a neighbour-labelling system. We added the human serum albumin signal peptide (Dugaiczyk, Law, and Dennison 1982), which is cleaved upon secretion, to create a secreted form of +36GFP which we refer to as s36GFP.

**Figure 1.**
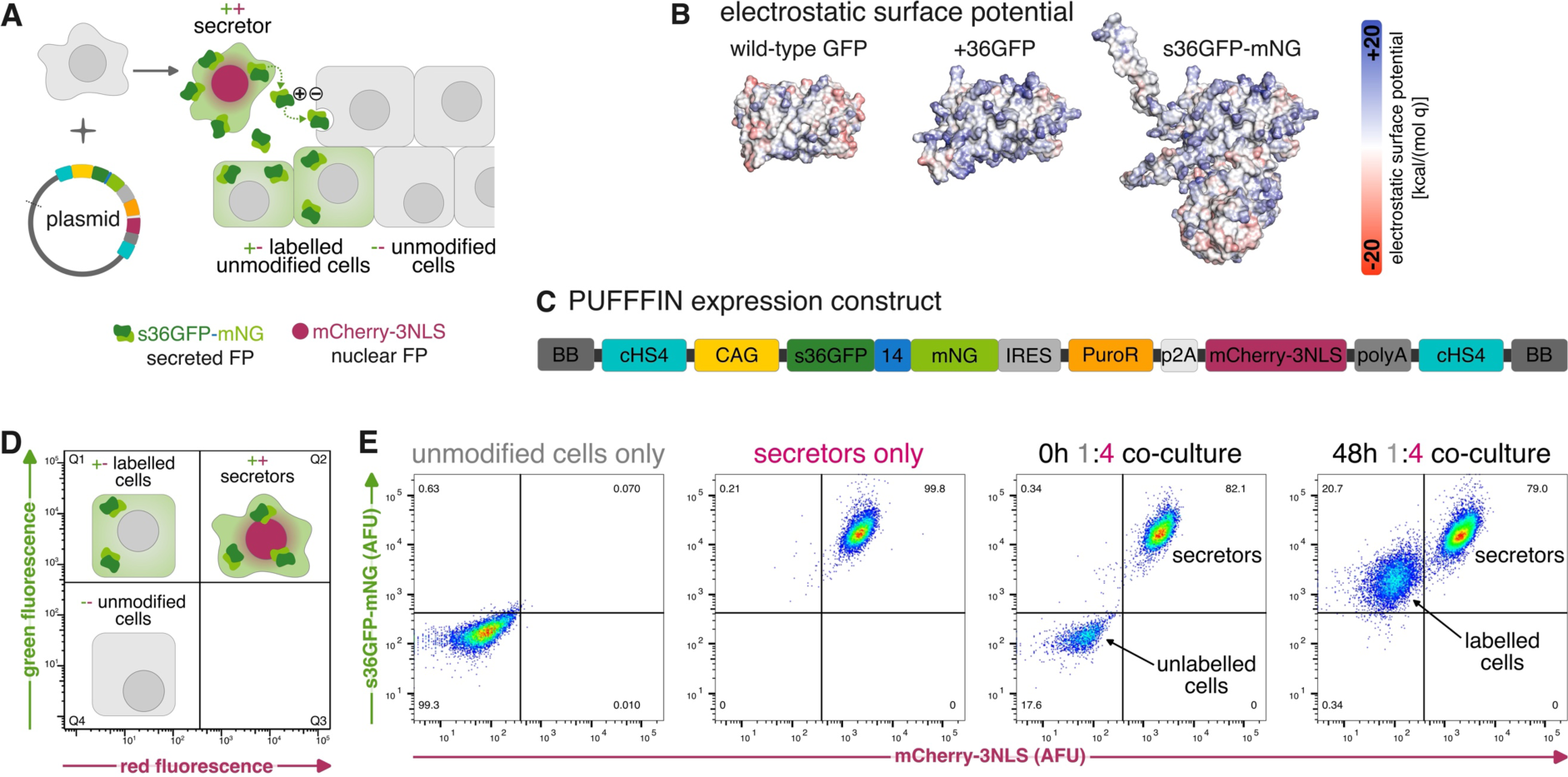
**(A)** PUFFFIN cell lines can be generated by transfection with the random integration plasmid containing the PUFFFIN expression construct. The resulting cells stably express s36GFP-mNG which is transferred to nearby cells while retaining nuclear mCherry-3NLS. **(B)** Electrostatic surface potential shown for wild-type GFP (PDB ID: 4kw4), +36GFP (AlphaFold predicted structure), and s36GFP-mNG (AlphaFold predicted structure). **(C)** The PUFFFIN expression cassette on the random integration plasmid contains the following parts: cHS4 insulators on both ends, the strong ubiquitous CAG promoter, the labelling protein s36GFP-mNG fused by a flexible 14 amino acid linker, an IRES for translation of downstream components, the puromycin resistance gene, a p2A self-cleaving peptide, nuclear mCherry-3NLS, and a synthetic polyA with a mammalian terminator. **(D)** Mock flow cytometry plot showing the three populations expected for PUFFFIN labelling. **(E)** Unmodified mCherry^-^ cells become s36GFP-mNG^+^ after mixing with mCherry^+^ s36GFP-mNG^+^ secretors in a 1:4 unmodified cell:secretor co-culture. 10,000 cells were analysed for the control panels and 30,000 cells for the 48h 1:4 co-culture. **Abbreviations:** +36GFP – green fluorescent protein with a theoretical net charge of +36; 14 – 14 AA linker; BB – backbone; CAG – CMV early enhancer/chicken β-actin promoter; cHS4 – chicken hypersensitive site 4 insulator; FP – fluorescent protein; GFP – green fluorescent protein; IRES – internal ribosome entry site; NLS – nuclear localisation signal; mNG – mNeonGreen; polyA – polyadenylation tail; PuroR – puromycin resistance gene; s36GFP – +36 green fluorescent protein with an N-terminal secretion signal.

In order to maximise the sensitivity, speed, and specificity of neighbour labelling we sought a strategy to boost fluorescence intensity. To achieve this, we fused s36GFP to the ultra-bright green-fluorescent mNeonGreen (mNG) (Shaner et al. 2013) as a signal amplifier (**Figure 1B**). This was the first step towards generating a system we call “Positive Ultra-bright Fluorescent Fusion For Identifying Neighbours“ (PUFFFIN).

The PUFFFIN plasmid also encodes mCherry-3NLS, which remains inside the nucleus and serves to distinguish secretors from surrounding cells that have taken up s36GFP-mNG (**Figure 1A, C, D**). The PUFFFIN expression cassette is driven by the strong ubiquitous CAG promoter (Niwa, Yamamura, and Miyazaki 1991; Dou et al. 2021) and cHS4 insulators flank the expression cassette to guard against silencing (Chung, Whiteley, and Felsenfeld 1993) (**Figure 1C**).

We delivered the PUFFFIN plasmid to HEK293 cells and generated stable monoclonal cell lines. The resulting “secretor” cell lines expressed mCherry-3NLS and s36GFP-mNG at levels that clearly distinguish secretors from unmodified cells by flow cytometry (**Figure 1E, Figure S1A**).

To test the neighbour-labelling capacity of s36GFP-mNG, we mixed a minority of unmodified cells with an excess of secretors (1:4 ratio) to ensure that unmodified cells were likely to be in proximity to secretors. Within 48h, over 95% of the unmodified cells became clearly s36GFP-mNG^+^ (**Figure 1E, Figure S1B**). This demonstrates that mammalian cells expressing the PUFFFIN plasmid unambiguously label surrounding cells.

### PUFFFIN labelling is confined to the local cell neighbourhood

A successful neighbour-labelling system depends on labelling being limited to nearby cells. To test this, we plated a small number of secretors among a large excess of unmodified cells (50:1 ratio) and performed time lapse imaging.

Over time, green fluorescence became associated with unmodified cells that were in the proximity of secretors, while more distant cells remained unlabelled (**Figure 2A, Video S1**). This indicates that s36GFP-mNG label is delivered only to nearby cells. To further assess the distance of labelling, we established a boundary between a secretor population and a population of unmodified cells, as illustrated in **Figure 2B**. The two populations were plated in two different wells separated by a removable two-well insert and cultured to confluency, upon which the insert was removed so both populations could expand towards each other.

**Figure 2.**
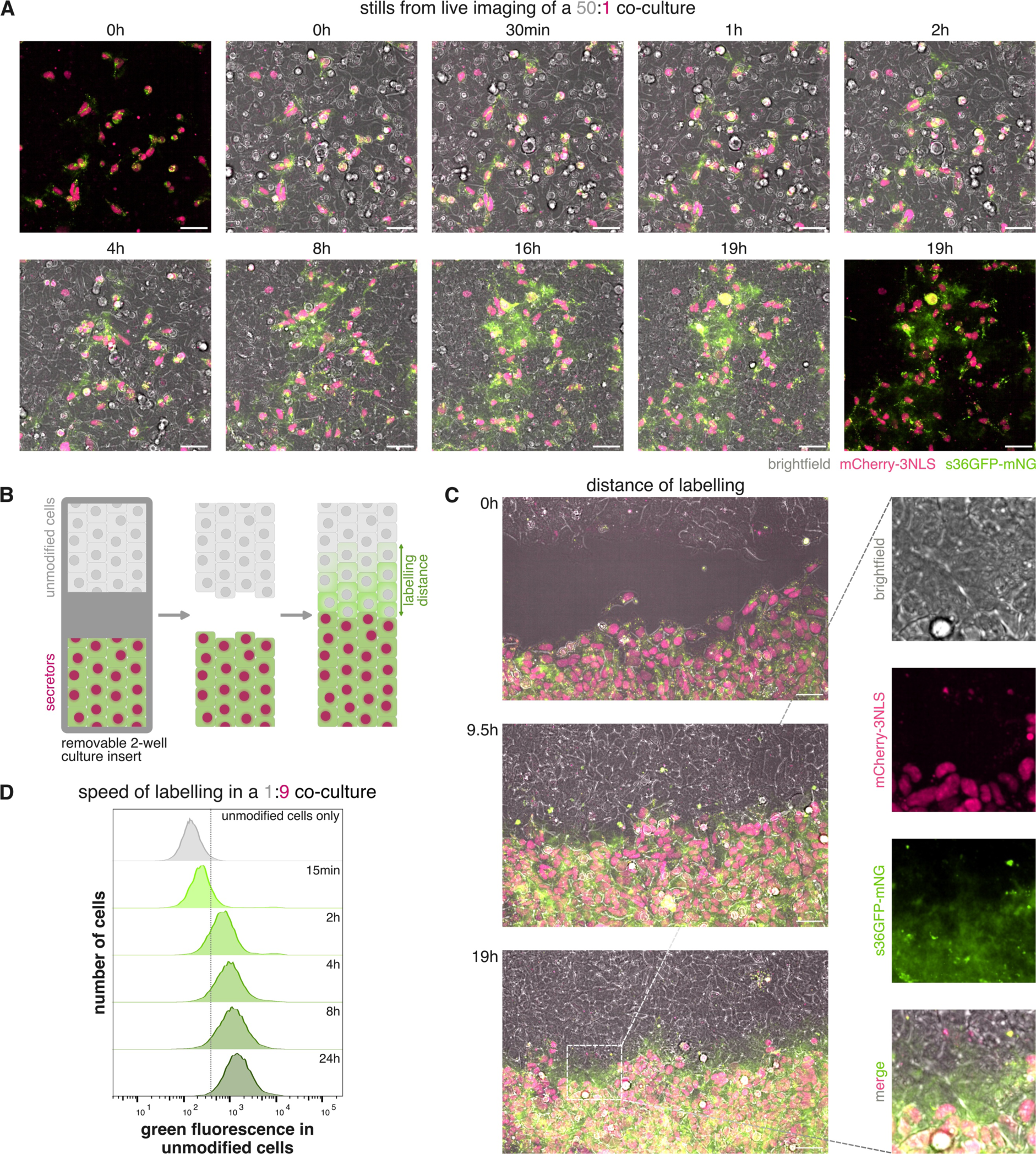
**(A)** Stills from live imaging of a 50:1 unmodified cell:secretor co-culture over 19 hours. Scale bars are 50 μm. **(B)** Schematic of experimental setup for testing the PUFFFIN labelling range. Secretors and acceptors are seeded separately and cultured to confluency in a removable cell culture insert. Insert is removed to allow both cell types to close the gap. Labelling at the border can be observed by fluorescence microscopy. **(C)** Stills from live imaging of a border experiment shows close labelling range. Scale bars are 50 μm. **(D)** PUFFFIN labelling is time dependent and fast: flow cytometry analysis of a 1:9 unmodified cell:secretor co-culture time course experiment. 10,000 unmodified cells were analysed for each sample.

s36GFP-mNG labelling became visible in unmodified cells that had reached the immediate neighbourhood of the secretor population, but was not detectable beyond four cell diameters from the boundary between the two populations in **Figure 2C and Video S2** (we note that this experiment is not designed to determine whether or not the green-fluorescent fusion protein can travel directly across several cell diameters, because the distance of labelling from the boundary will be dependent on cell division and cell migration as well as direct transfer of label).

Taken together, these data confirm that detectable label is delivered only to cells that are in close proximity to secretors.

### PUFFFIN labelling is transferred rapidly between cells

During the time-lapse experiments described above (**Figure 2A, C**) we could detect green fluorescence within neighbours of secretors within 30 minutes. To further assess how long it takes for the s36GFP-mNG label to become detectable, we plated unmodified cells with an excess of secretors (1:9 ratio) at high density and then analysed green fluorescence within the mCherry-negative unmodified cell fraction at various time points. s36GFP-mNG was detected in unmodified cells within 15 minutes of mixing them with PUFFFIN secretors. Labelling intensity increased further after 2h of co-culture and continued to increase over the next 22h (**Figure 2D**).

We conclude that PUFFIN labelling is transferred to neighbouring cells within minutes at levels that can be detected by flow cytometry or by live imaging.

### The PUFFFIN system has a modular design to combine manipulation of cell function with customisable neighbour labelling within a single plasmid

To achieve ready customisability, we use the Extensible Mammalian Modular Assembly (EMMA) platform (Martella et al. 2017) to combine all components of PUFFFIN within a single plasmid comprised of modular exchangeable parts (**Figure 3**).

**Figure 3.**
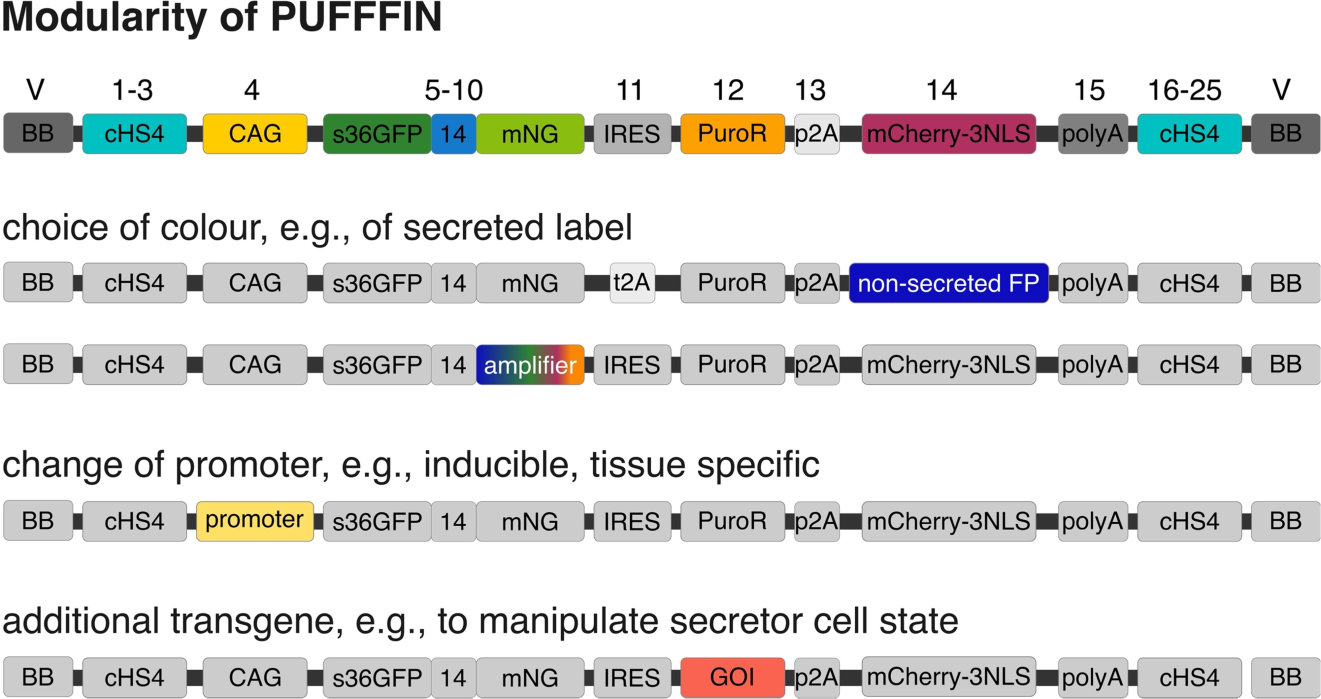
Modular design of PUFFFIN plasmid assembly. Assembly of different parts allows for example combination of the s36GFP-mNG label with a non-secreted fluorescent protein (FP) in another colour. The PUFFFIN label could be a fusion of s36GFP with an amplifier other than mNeonGreen to change the colour of the label. The ubiquitous CAG promoter could be replaced by a cell type specific promoter and the selection gene replaced by a gene of interest.

For example, to make a version of PUFFFIN that is compatible with cells that already contain red-fluorescent reporters, we exchanged the mCherry-3NLS transgenes to a cytoplasmic blue-fluorescent tagBFP (Subach et al. 2008), and at the same time replaced the IRES with a t2A self-cleaving peptide (Liu et al. 2017). We used this customised construct to generate a stable polyclonal line of secretors in which the vast majority of cells (> 95%) are s36GFP-mNeonGreen^+^ tagBFP^+^ (**Figure S2**), without the need for selecting clonal cell lines.

To expand the utility of the system for functional experiments, the puromycin resistance gene can be exchanged for any transgene of interest for manipulating or monitoring cell behaviour (the selection gene is dispensable because stably expressing secretors can be purified based on fluorescent signal rather than antibiotic selection). Similarly, the CAG promoter can be switched to an inducible or a cell-type-specific promoter for regulatable or tissue-specific expression of the PUFFFIN construct. This offers the opportunity to manipulate cell behaviour in a specific set of cells while simultaneously delivering a fluorescent label to their neighbours.

### The PUFFFIN system can be customised for colour-of-choice labelling using HaloTag® technology

We demonstrate above (**Figure 1 and Figure 2**) that mNeonGreen is an effective signal amplifier that facilitates bright neighbour labelling using PUFFFIN. However, s36GFP-mNeonGreen labelling is not compatible with cells that already express GFP, such as green-fluorescent reporter lines. We therefore sought an approach to further customise the PUFFFIN construct so that labelling can be performed in any colour of choice without the need for additional genetic manipulation to the PUFFFIN plasmid.

To achieve this, we replaced the coding sequence of mNeonGreen with a HaloTag® (Los et al. 2008), illustrated in **Figure 4A, B, Figure S3A**, creating a s36GFP-HT fusion protein (‘PUFFHalo’) that can bind any HaloTag fluorescent ligand (England, Luo, and Cai 2015; Cook, Walterspiel, and Deo 2023) (**Figure 4C**). We used this new construct to generate stable PUFFHalo secretor lines.

**Figure 4.**
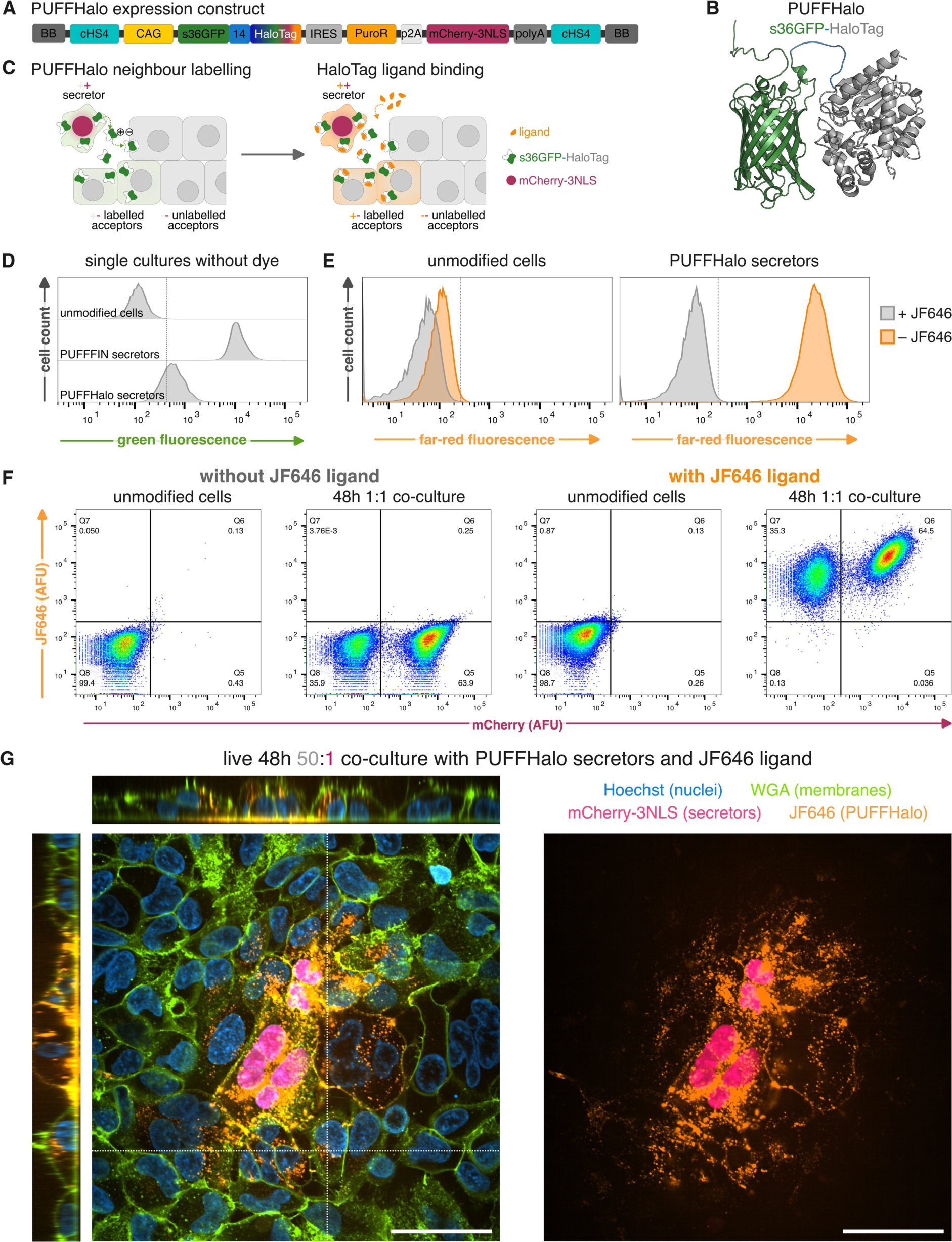
**(A)** The modularity of PUFFFIN allows an assembly termed “PUFFHalo” that contains HaloTag as amplifier instead of mNeonGreen. **(B)** As an alternative to the s36GFP-mNG label, s36GFP (green) was fused to HaloTag (grey) as signal amplifier by a 14 amino acid-long linker (blue) (AlphaFold predicted structure). **(C)** Neighbour labelling is possible with PUFFHalo but has only low green fluorescence on its own. The label can be bound by HaloTag ligands, e.g., dyes in different colours, which amplifies the labelling signal. **(D)** Comparison of unmodified cells, s36GFP-mNG secretors and PUFFHalo secretors shows low green-fluorescence of the unstained PUFFHalo. **(E)** Unmodified cells and PUFFHalo secretors were incubated for 2h with or without the HaloTag ligand Janelia Fluor (JF646). **(F)** Flow cytometry of 48h 1:1 co-cultures of unmodified cells and PUFFHalo secretors were incubated for 2h with or without the HaloTag ligand JF646. **(G)** Live imaging of a 48h 50:1 co-culture of unmodified cells and PUFFHalo secretors with the HaloTag ligand JF646, with Hoechst nuclear staining and WGA488 membrane staining. Dotted lines mark regions selected for orthogonal views. Left image shows merge of all channels, right only far-red JF646 (PUFFHalo) and red (mCherry-3NLS, secretor nuclei). Scale bars are 50 μm, image depth is 11.5 μm.

We first confirmed that the residual green fluorescence from the s36GFP was relatively dim in the absence of any amplifier, and therefore likely to be compatible with GFP-based reporters, and unlikely to override our ability to customise the colour of PUFFFIN labelling using HaloTag-binding dyes **(Figure 4D)**. We then set out to test whether we could customise the colour of PUFFFIN labelling using fluorescent HaloTag ligands. We cultured secretors in the presence or absence of far-red (JF646) or green (OregonGreen) fluorescent dyes and observed a significant shift in fluorescence following dye addition (**Figure 4E, Figure S3B**). This suggests that PUFFHalo can be labelled with different fluorophores and detected by flow cytometry.

We next tested the ability of PUFFHalo to label neighbours, by mixing secretors with unmodified cells in a 1:1 co-culture at high density for 48h hours. More than 99% or 85% unmodified cells were labelled by the PUFFHalo in the presence of far-red or green HaloTag fluorescent ligands, respectively (**Figure 4F, Figure S3C**). This suggests PUFFHalo can be transferred to unmodified neighbours of secretor cells, and that labelled neighbours can be clearly identified by flow cytometry.

Finally, we investigated the potential of PUFFHalo for live imaging applications, by mixing a low number of secretors with an excess of unmodified cells (50:1 ratio) for 48h and imaging the cells following addition of HaloTag dyes. Labelling was clearly visible within neighbours of mCherry^+^ secretors and could also be seen decorating the cell membrane and underlying substrate (**Figure 4G, Figure S3D, E**). This suggests that PUFFHalo can be used to monitor neighbour-labelling in live cells in real-time.

We conclude that PUFFHalo is readily customisable using Halo-compatible dyes to offer labelling in a colour of choice, and that the Halo-compatible dyes can further enhance the sensitivity of neighbour labelling.

## Conclusion

We demonstrate that Positive Ultra-bright Fluorescent Fusion For Identifying Neighbours (PUFFFIN) can be used to unambiguously, sensitively, and rapidly label the neighbours of mammalian cells within minutes. Our modular design can incorporate any transgene of interest to study non-cell-autonomous effects of experimentally induced changes in identity of behaviour. The integration of PUFFFIN with HaloTag technology, PUFFHalo, enables colour-of-choice labelling with high signal-to-noise ratio without the need to reengineer the PUFFHalo plasmid. The HaloTag could in principle also be used for supporting extended live imaging (Cook, Walterspiel, and Deo 2023), pulse-chase experiments (Yim, Yamamoto, and Mizushima 2022), and specific degradation of the label using HaloPROTACS (Buckley et al. 2015).

PUFFFIN offers a high level of customisability, a convenient single-plasmid delivery system, and receptor-independent transfer that does not require any modifications to the cells that receive the label. This makes PUFFFIN user-friendly and readily adaptable, offering an effective and flexible tool for illuminating cellular neighbourhoods.

## Materials and Methods

### Construct design

We chose the following parts for the PUFFFIN construct design: the strong, ubiquitous CAG promoter (Niwa, Yamamura, and Miyazaki 1991) should ensure high expression levels of the PUFFFIN construct and is followed by a Kozak consensus sequence (GCCGCCACC). The PUFFFIN fusion proteins start with a secretion signal peptide (s) from the human serum albumin (HSA) precursor (Dugaiczyk, Law, and Dennison 1982), optimised for mammalian expression by balanced GC content and improved codon adaptation index (0.96 instead 0.65 in original sequence), followed by a short 2 amino acid RG linker to maintain the sequence context of the HSA precursor, to prevent issues with cleavage of the signal peptide upon secretion. The sequence of +36GFP (Lawrence, Phillips, and Liu 2007) is codon optimised for expression in mammalian cells (Bar-Shir et al. 2015), the N-terminal His-tag was removed from the original sequence. The 14 amino acid-long linker is glycine-serine rich for stability while the small polar amino acid threonine makes it flexible (Chen, Zaro, and Shen 2013). To increase the fluorescent signal of s36GFP, it was fused to an amplifier: either the ultra-bright green-fluorescent protein mNeonGreen (Shaner et al. 2013) or a HaloTag (Los et al. 2008). An internal ribosomal entry site (IRES) follows the s36GFP-amplifier coding sequence, to allow expression of a downstream puromycin N-acetyltransferase (PAC) gene, which confers resistance (PuroR) to the antibiotic puromycin dihydrochloride (Luna et al. 1988), enabling selection of clones with stable integration of the PUFFFIN constructs. The resistance gene can be replaced with a functional transgene of interest (GOI). A 2A self-cleaving peptide from porcine teschovirus-1 (p2A) (X. Tang et al. 2016) separates PAC and the subsequent fluorescent protein. The red-fluorescent mCherry (Shaner et al. 2004) was fused to three C-terminal SV40 nuclear localisation signals (3NLS) to drive nuclear localisation (Malaguti et al. 2013; 2022). Alternatively, cytoplasmic blue-fluorescent tagBFP (Subach et al. 2008) was used instead of mCherry-3NLS. A STOP cassette, consisting of a synthetic poly-A (SPA) site (Levitt et al. 1989) and a C2MAZ terminator-binding sequence (Yonaha and Proudfoot 2000) halts downstream transcription and translation. The whole expression construct is flanked by chicken β-globin insulators (cHS4) to prevent silencing at the random integration site (Chung, Whiteley, and Felsenfeld 1993). This was confirmed by culturing secretors to passage 15 without a decrease in s36GFP-mNG or mCherry-3NLS expression levels determined by flow cytometry. Ampicillin can be used for selective outgrowth of competent cells. We recommend linearisation with PvuI restriction enzyme.

### Protein folding and charge prediction

Protein folding prediction for +36GFP, s36GFP-mNG, and s36GFP-HT was done with ColabFold (Mirdita et al. 2022). Bluues Version 2.0 (Walsh et al. 2012) was used for prediction of theoretical net charge and electrostatic surface potential of the folded sequences and wild-type GFP (PDB ID: 4kw4).

### EMMA cloning: DNA domestication, assembly, and purification

In the original EMMA toolkit publication the authors domesticate DNA parts by subcloning them into different entry vectors for each position in the assembly (Martella et al. 2017). We modified this strategy in order to use the same entry vector for each position. PCR primers for DNA domestication were designed as follows: 5’-(N)_6_-BsmBI binding site-N-fusion site-(N)_20_-3’. As an example, the primers for position 1 were: Forward: 5’-(N)_6_-CGTCTC-N-TAGG-(N)_20_-3’, Reverse: 5’-(N)_6_-CGTCTC-N-CCAT-(N)_20_-3’. PCR amplification was performed with a Q5 HotStart High Fidelity Polymerase (NEB, #M0493S), and resulting blunt-ended PCR products were cloned into a pCR-Blunt II-TOPO vector (Invitrogen Zero Blunt TOPO PCR Cloning Kit; #450245) to generate domesticated DNA. This strategy allows for the creation of functional DNA sequences spanning multiple positions, without the need for assembly connectors. New DNA sequences, e.g., for the s36GFP-mNG fusion protein, were custom-synthesised by GeneArt or Integrated DNA Technologies.

TOPO Cloning reactions were transformed into NEB® 10-beta Competent E. coli (High Efficiency) (NEB, #C3019I) following manufacturer’s instructions. Bacteria were plated onto pre-warmed kanamycin resistance LB agar plates in the presence of X-gal (Promega, #V3941) for blue/white screening, and incubated overnight at 37 °C. White colonies were picked into 5 ml LB in the presence of kanamycin, and incubated at 37 °C prior to DNA purification.

EMMA assemblies of parts into the YCe3736_HC_Amp_ccdB_receiver vector (a kind gift from Prof Steven Pollard, also available on Addgene: #100637) were carried out using the NEBridge Golden Gate Assembly Kit (BsmBI-v2) (NEB, #E1602L), using 75 ng of each part and 2 μl of enzyme mix in a 20 μl reaction, performing 60 cycles of digestion (42 °C, 5 min) and ligation (16 °C, 5 min), followed by heat inactivation of enzymes (60 °C, 5 min). For assemblies involving domesticated parts harbouring extra BsmBI restriction sites, an extra ligation step was performed following heat inactivation: the reaction volume was increased to 30 μl, adding 400 U (1 μl) T4 DNA Ligase (NEB, #M0202S), and incubating the reactions at 16 °C for an additional 5-16 h.

10 μl of the assembly reaction was transformed into NEB® 5-alpha Competent E. coli (Subcloning Efficiency) (NEB, #C2988J) following manufacturer’s instructions. Bacteria were plated onto pre-warmed ampicillin resistance LB agar plates at two densities: 1/10 and 9/10 of the transformation reaction, and incubated overnight at 37 °C. Colonies were picked into 5 ml LB in the presence of ampicillin, and incubated at 37 °C prior to DNA purification. In our experience, colonies obtained with subcloning efficiency E. coli usually contain the correctly assembled plasmid. Should no colonies be obtained with subcloning efficiency E. coli, the remaining 10 μl of the assembly reaction can be transformed into high efficiency E. coli. In our experience, in this instance it is necessary to screen more bacterial colonies to identify correct assemblies.

Plasmid DNA was purified using the Monarch Plasmid DNA Miniprep Kit (NEB, #T1010), and subjected to diagnostic restriction digest and subsequent gel electrophoresis. Sequences of clones with correct banding patterns were verified by either Sanger sequencing of inserts or whole plasmid sequencing with Oxford Nanopore Technologies.

### Cell line generation and maintenance

HEK293 cells were transfected with 3 μg of linearised plasmid using Lipofectamine 3000 Transfection Reagent (Invitrogen, #L3000001) in wells of a 6-well plate. Selection with 2 μg/ml puromycin dihydrochloride (Sigma-Aldrich; #P8833) was started 48h after transfection for transfected cells and a mock transfection control. Cells were replated from the 6-well plate at 1/10 and 9/10 densities onto 10 cm cell culture dishes and resulting colonies picked and expanded as monoclonal lines. Polyclonal lines were considered to only contain stable integrants following death of all cells in the mock transfection control dishes. All cell lines were maintained in HEK293 culture medium – Dulbecco’s Modified Eagle Medium (DMEM, high glucose, pyruvate; Gibco, #41966029) with 10% foetal calf serum (FCS; Biosera, #FB-1280/500) – under standard culture conditions (37 °C and 5% CO_2_) and passaged after washing with Dulbecco’s phosphate buffered saline (PBS; Sigma-Aldrich, #D8537) using 0.025% trypsin (Invitrogen, #15090-046). To enrich the stable polyclonal s36GFP-HT secretor line for high transgene expressing cells, cells were prepared for flow cytometry as described below, and sorted based on fluorescent marker expression on a BD FACSAria II. After sorting, cells were cultured for two weeks in culture medium supplemented with penicillin/streptomycin (100 U/ml; Invitrogen, #15140-122).

### Co-culture experiments

Secretors and unmodified cells were detached from cell culture vessel using trypsin, resuspended in culture medium, pelleted by centrifuging at 300 g for 3 min, resuspended in an appropriate amount of medium and counted to plate cells at unmodified cells-to-secretor ratios indicated for each experiment. Single cultures for secretors and acceptors were included for all experiments.

### Flow cytometry

For flow cytometry experiments, cells were cultured in 12-well plates for 2h – 48h time points, mixed as cell suspensions for 15 min time points, or mixed on ice immediately prior to analysis for 0h time points. Cells were detached using trypsin, quenched in HEK293 culture medium, pelleted by centrifuging at 300 g for 3 min, and resuspended in 10% FCS in PBS with either DAPI (1 μM; Biotium, #40043) or DRAQ7 (300 nM; Abcam, #ab109202) live/dead staining. No DAPI/No DRAQ7 controls were included for all experiments. Samples were analysed on a BD LSRFortessa Cell Analyzer using V 450/50-A (blue fluorescence), B 530/30-A (green fluorescence), Y/G 610/20-A (red fluorescence), and R 670/14-A (far-red fluorescence) laser/filters combinations.

### Live imaging

Cells were plated on 7.5 μg/ml fibronectin-coated (Sigma-Aldrich, #F1141) 4-well μ-slides (Ibidi, #80426) with or without removable culture-inserts (Ibidi, 2-well #80209 or 3-well #80369). Culture inserts were removed 8h prior to imaging and HEK293 culture medium was changed to imaging medium – phenol red-free DMEM (Gibco, #31053028) with 10% FCS (Biosera, #FB-1280/500) and 2% glutamate/pyruvate (100mM sodium pyruvate, Invitrogen #11360-039; 200mM L-glutamine, Invitrogen, #25030-024). Slides were imaged for brightfield, red and green fluorescence at 15 min intervals over 19h on an Opera Phenix™ High-Content Screening System (PerkinElmer) under standard culture conditions.

### HaloTag staining and analysis

Cells were plated for co-culture experiments as described above, either in 12-well plates for flow cytometry or in fibronectin-coated 8-well μ-slides for live imaging. On the day of analysis, HEK293 culture medium was replaced by imaging medium containing either Janelia Fluor 646 (JF646) ligand (10 nM; Promega, #GA1120) or OregonGreen ligand (50 nM; Promega, #G2801) or no HaloTag dye. Cells were incubated for 2h at 37 °C and prepared for and processed by flow cytometry as described above. Hoechst 33342 (100 ng/ml; Apexbio, #A3472-APE) nuclear staining was added to the medium of the 8-well slide 1h prior to imaging, and WGA488 (1 μg/ml; Wheat Germ Agglutinin CF®488A Conjugate; Biotium, #BT29022-1) membrane staining was added to single wells immediately before imaging on an Opera Phenix™ High-Content Screening System (PerkinElmer).

## Funding

T.L. was supported by the Integrative Cell Mechanisms PhD Programme, with funding by Wellcome [218470], and the University of Edinburgh School of Biological Sciences and a core grant to the Wellcome Centre for Cell Biology [203149].

M.M. is supported by a BBSRC Engineering Biology Transition Award (BB/W014610/1), and by a University of Edinburgh School of Biological Sciences internal Seeding Fund award.

S.L. is supported by a Wellcome Trust Senior Fellowship [220298]

## Supporting information

Video S1 Lebek

Video S2 Lebek

## Acknowledgments

We would like to thank Mihaly Badonyi for support with construct design of the s36GFP label and the s36GFP-mNG and PUFFHalo fusion proteins. We thank Charles AC Williams for help with HaloTag staining and imaging support. We are thankful to Rachel White and Steven M Pollard for providing the YCe3736_HC_Amp_ccdB receiver vector and assistance with EMMA cloning. Thank you to the group members of the Lowell Lab, especially but without any particular order, to Jen Annoh, Maria Rosa Portero Migueles, Eleanor Earp, Aisling Fairweather, Alicia Louis Perez Lezcano, and Matthew French. We thank the Wilson and Blin Labs for valuable feedback and reagents shared. We are grateful to the facilities at the Institute for Regeneration and Repair of the University of Edinburgh, especially Fiona Rossi at the Flow Cytometry Core Facility, Justyna Cholewa-Waclaw at the High Content Screening Facility, and Theresa O’Connor at the Tissue Culture Facility.

## Supplementary material

### Supplementary Figures

**Figure S1.**
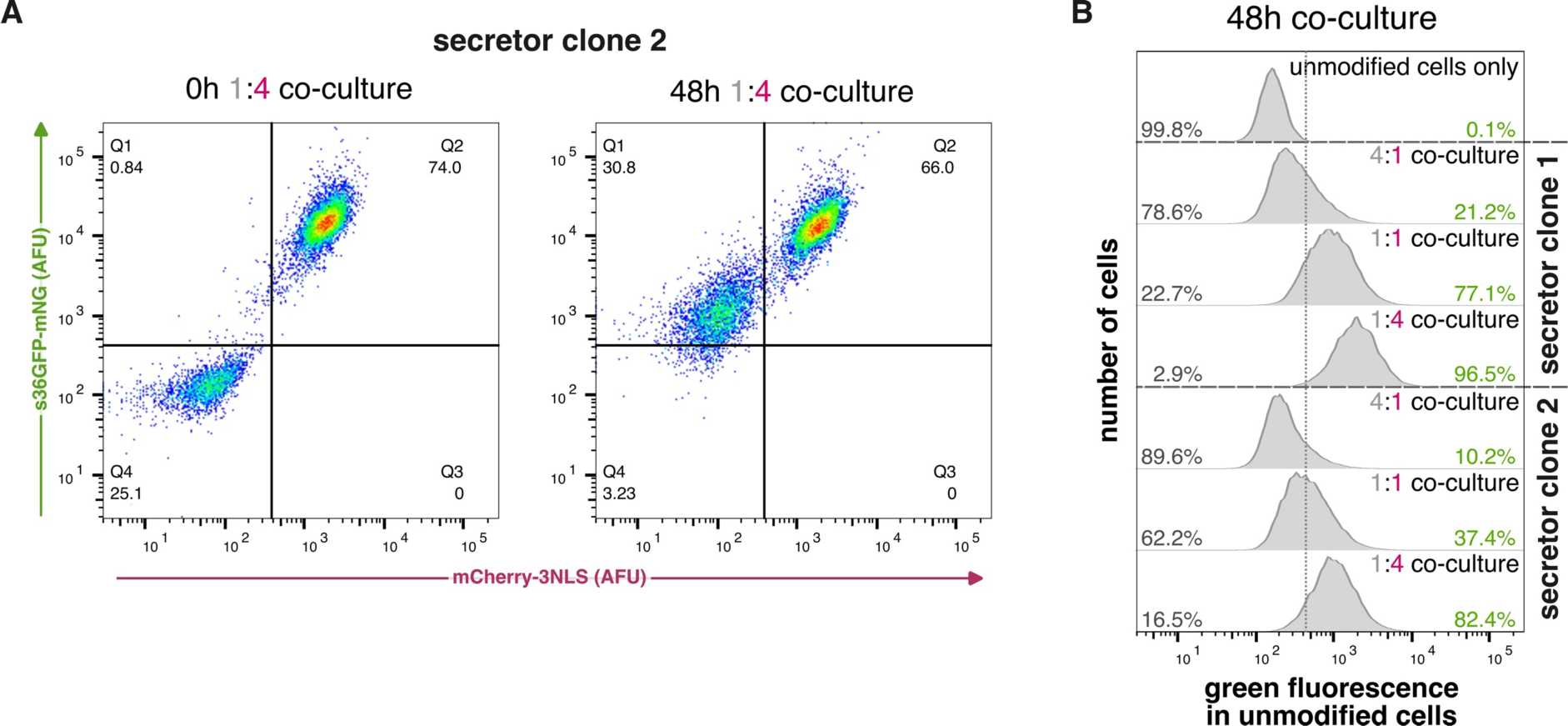
A second independent monoclonal secretor line can effectively label unmodified cells by transferring s36GFP-mNG. **(A)** Flow cytometry of a 1:4 co-culture of unmodified cells and secretor clone 2 at 0h and 48h time points. **(B)** Flow cytometry of a 48h co-culture with unmodified cells and either secretor clone 1 or secretor clone 2 seeded at different co-culture ratios. Grey numbers are percentage of acceptors in Q4, green numbers are percentage of acceptors in Q1.

**Figure S2.**
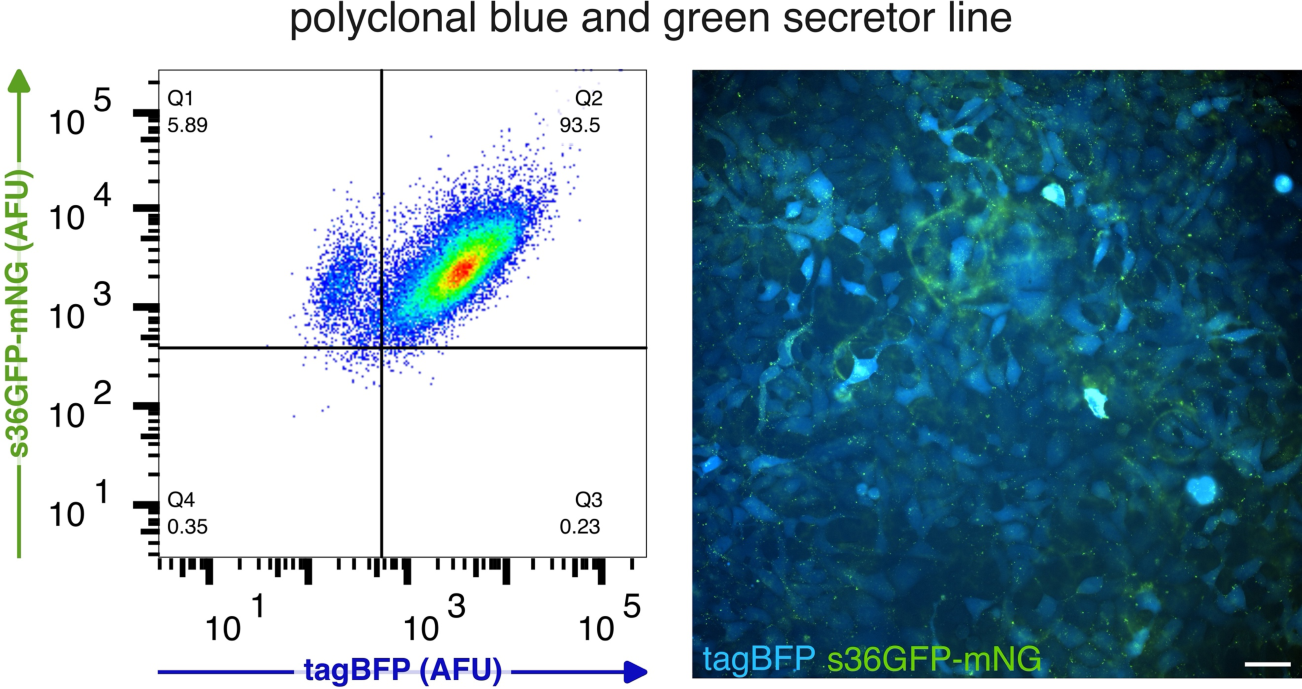
Modularity of the PUFFFIN expression construct allows change of colour for the secretor-cell-marker. Polyclonal secretors expressing s36GFP-mNG and cytoplasmic tagBFP as shown by flow cytometry (left) and imaging live cells for green and blue fluorescence (right). Scale bar is 50 μm.

**Figure S3.**
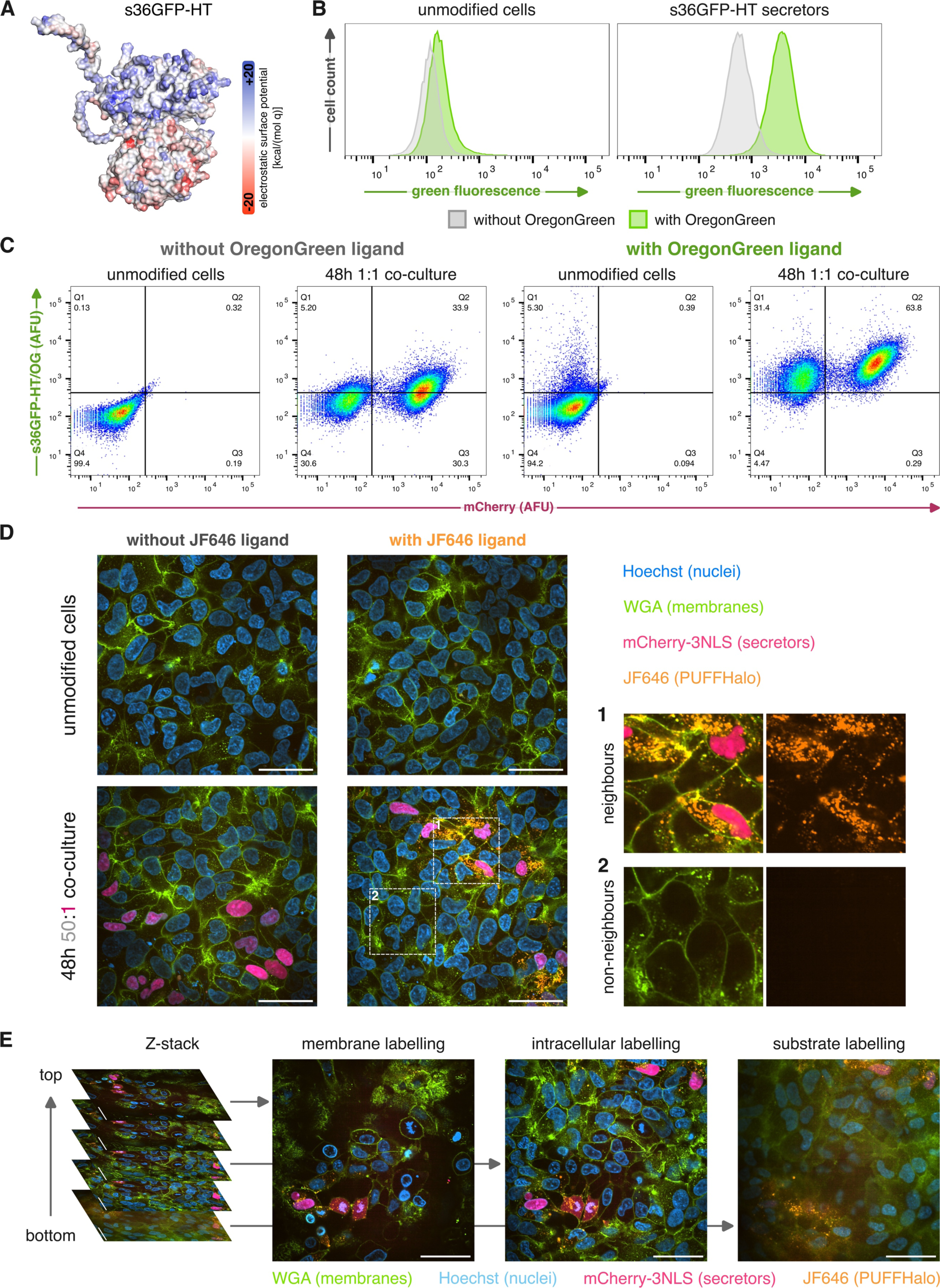
PUFFHalo integrates HaloTag technology with PUFFFIN for choice-of-colour labelling. **(A)** Electrostatic surface potential shown for s36GFP-HaloTag (AlphaFold predicted structure). **(B)** Unmodified cells and PUFFHalo secretors were incubated for 2h with or without the HaloTag ligand OregonGreen. **(C)** Flow cytometry of 48h 1:1 co-cultures of unmodified cells and PUFFHalo secretors were incubated for 2h with or without the HaloTag ligand OregonGreen. **(D)** Live imaging of single cultures of unmodified cells and 48h 50:1 co-cultures of unmodified cells and PUFFHalo secretors with or without the HaloTag ligand JF646, all with Hoechst nuclear staining and WGA488 membrane staining. Two regions of the 48h 1:1 co-culture with JF646 are magnified to show neighbours and non-neighbours. Scale bars are 50 μm. **(E)** Three single planes of a Z-stack for a 48h 1:1 co-culture with JF646, WGA488, and Hoechst staining were selected to highlight different localisations of the s36GFP-HT label. Scale bars are 50 μm.

## Notes

### Competing Interest Statement

The authors have declared no competing interest.

### Summary of Updates

Correction to typo in title of paper

